# The P3018S disease variant reveals how dynein’s trailing motor sets ensemble velocity

**DOI:** 10.64898/2026.05.14.725143

**Authors:** Caitlyn A. Walker, Lu Rao, Abhipsa Shatarupa, Xinglei Liu, Jun Yang, Hernando Sosa, Kai Zhang, Arne Gennerich

**Affiliations:** Department of Biochemistry and Gruss Lipper Biophotonics Center, Albert Einstein College of Medicine, Bronx, NY 10461, USA; Department of Molecular Biophysics and Biochemistry, Yale University, New Haven, CT 06511, USA

## Abstract

Mutations in the cytoplasmic dynein heavy chain (DYNC1H1) underlie a range of neurodevelopmental disorders yet how individual variants perturb dynein function remains poorly understood. We characterize the disease-associated P3018S mutation, located in the AAA4 module and within the Lis1-interacting region of the motor. Dynein–dynactin–BicD2 (DDB) complexes containing P3018S dynein move at half the wild-type velocity and generate reduced stall forces but retain the ability to assume the inhibitory phi conformation that Lis1 suppresses, respond to Lis1, and assemble into higher-order force-generating states. Several *Schizosaccharomyces* species, which lack Lis1, naturally encode a serine at this position, suggesting evolutionary relevance for dynein activation. Using mixed wild-type–mutant assemblies, we find that the dynein occupying the trailing position on dynactin dictates ensemble velocity, and that the leading dynein’s LIC enhances trailing-motor velocity by ∼130%, revealing a mechanism for leading-to-trailing motor stimulation within multi-dynein assemblies.

## INTRODUCION

Cytoplasmic dynein-1 (hereafter “dynein”) is the primary minus-end-directed microtubule motor in eukaryotic cells. It transports diverse cargoes—including organelles, proteins, mRNAs, and viruses^1–3^—over long cellular distances, and disruptions in dynein function cause a growing set of human neurological disorders collectively termed “dyneinopathies”^4–6^, including spinal muscular atrophy (SMA)^7,8^, SMA with lower-extremity predominance (SMALED)^9–12^, and malformations of cortical development^13–17^. We recently learned of a heterozygous missense mutation in DYNC1H1, c.9052C>T (p.P3018S), identified clinically in a boy treated at the Einstein–Montefiore Medical Center^18^. The variant, which is absent from population databases, is classified as likely pathogenic. The patient presented with global developmental delay beginning around one year of age and continues to exhibit impairments in motor planning, coordination, learning, and attention at age 15. Because dynein is essential for neuronal development and intracellular transport, defining how the P3018S mutation alters motor function is essential for understanding its potential contribution to the patient’s phenotype.

Activation of mammalian dynein requires binding to its largest cofactor, dynactin, together with a coiled-coil cargo adaptor such as BicD2^19,20^. Dynein on its own is diffusive^21^ or weakly processive^41,42^, but formation of the tripartite dynein–dynactin–adaptor complex converts it into an highly processive motor (the DDB complex)^19,20,22^. This activation involves a transition from the autoinhibited “inverted” or phi-like conformation^23–31^ (named for its resemblance of the Greek letter Φ) to an open, parallel arrangement of the motor domains^25,27,28,32^ (**Fig. 1a**). Once activated, dynein undergoes coordinated stepping under load^33–35^, with one motor domain tightly engaged with the microtubule while the other advances. Structural and biochemical studies further show that dynactin–adaptor complexes can recruit two dynein dimers, positioning them in a staggered, parallel configuration that supports cooperative motility and enhanced force generation^33,36^ (**Fig. 1a**).

**Figure 1.**
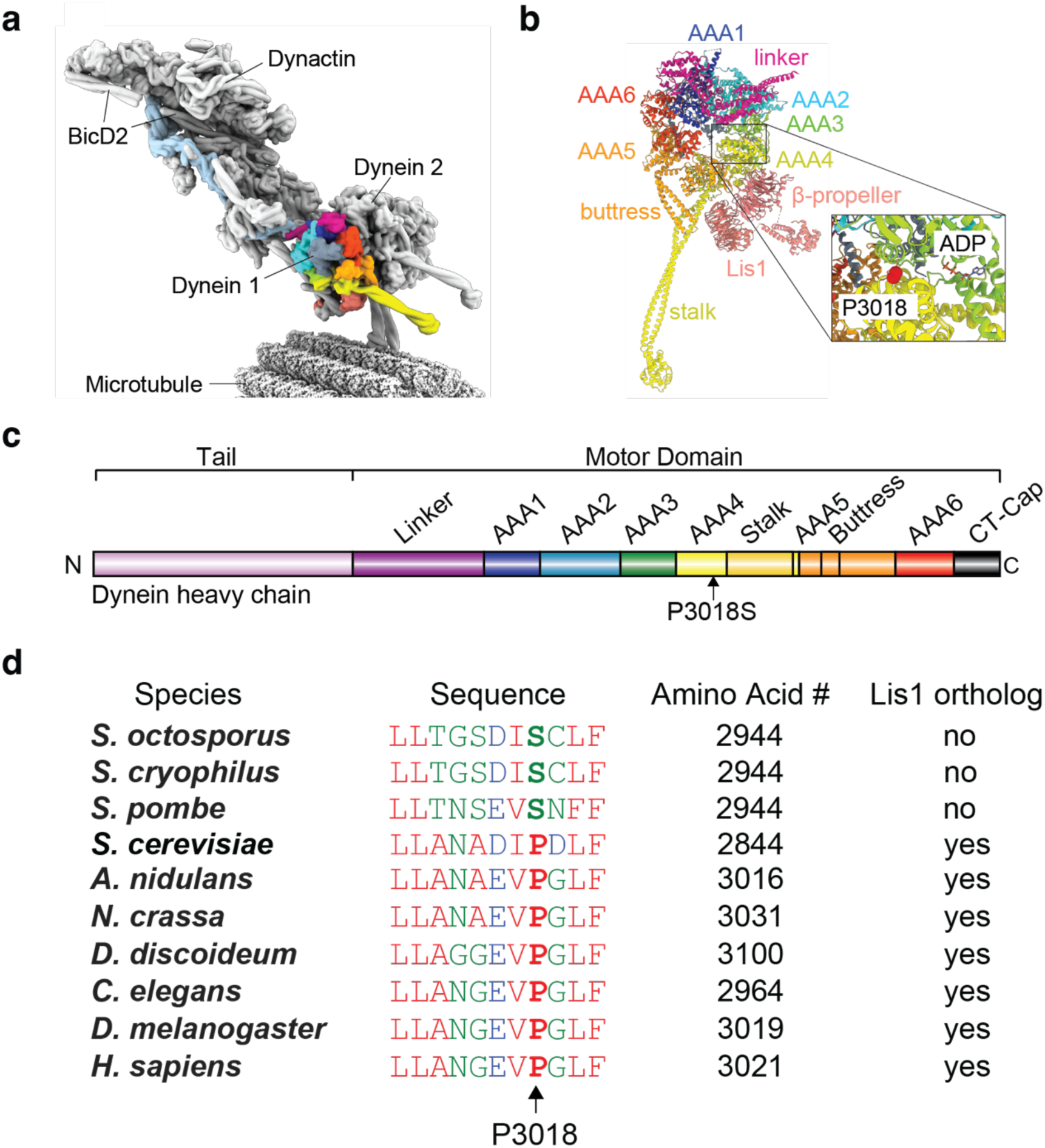
Structural context of the P3018 residue in Lis1-bound dynein within the DDJ complex. **(a)** CryoEM structure of the dynein-dynactin-JIP3 (DDJ) complex containing two dyneins, one JIP3 dimer, dynactin, and Lis1 bound to a microtubule (PDB 8PTK). The two dynein motor domains adopt a parallel arrangement along the microtubule. One dynein motor is shown with individual AAA+ domains colored as follows: AAA1, blue; AAA2, cyan; AAA3, green; AAA4, yellow; AAA5, orange; AAA6, magenta; C-terminal cap, gray. The tail is shown in light blue. **(b)** Enlarged view of a single dynein motor domain highlighting Lis1’s two β-propellers bound to AAA4 and the base of the stalk. Color coding is as in (a). The inset indicates the position of residue P3018 within the AAA4 domain. **(c)** Schematic representation of the domain organization of the dynein heavy chain, including the tail, linker, AAA+ ring, stalk, and microtubule-binding domain. **(d)** Sequence alignment of dynein heavy chains from selected species showing that organisms with a naturally encoded serine at the position corresponding to human P3018 lack an identifiable Lis1 ortholog.

A key regulator of dynein activation is Lis1, a dynein-binding cofactor whose dysfunction causes Type I lissencephaly^37–41^. Lis1 binds AAA4 and the base of the stalk (**Fig. 1b**), stabilizing dynein in an open conformation and preventing formation of the autoinhibited phi state^23–31^, thereby promoting assembly of two-dynein dynein–dynactin–adaptor (DDA) complexes^42,43^, the fully activated form of the motor. Because Lis1 dissociates once motors engage microtubules^44^, it is thought to function primarily as a priming factor rather than as a regulator of dynein’s movement.

Our recent biophysical studies demonstrated that the phi-like conformation limits the maximal force that a single dynein in a DDB complex can generate to ∼2.5 pN, whereas preventing phi formation—through Lis1 binding^23,44,45^ or mutations that block the phi state^25^—allows a single dynein to reach ∼4.5 pN. When two-dynein DDB complexes are formed, their motors can jointly sustain ∼7 pN at stalling. Under higher load, these two-dynein assemblies can recruit a third dynein through an auxiliary adaptor, yielding three-dynein DDB complexes capable of generating ∼9 pN. Together, these observations reveal a load-dependent reorganization of dynein from one- to two-to three-motor states, each supporting progressively greater force output.

Here we define how the P3018S mutation alters dynein’s mechanochemistry, regulation by Lis1, and cooperative motility within dynein–dynactin–BicD2 (DDB) assemblies. We find that P3018S dynein moves at approximately half the velocity of wild-type (WT) dynein and generates reduced stall forces, yet it retains the ability to form the autoinhibited phi conformation, respond to Lis1, and assemble into higher-order force-generating complexes. Remarkably, several *Schizosaccharomyces* species—which lack Lis1—naturally encode a serine at this position, raising the possibility that P3018S may influence dynein’s activation pathway. Using mixed WT–mutant DDB complexes, we show that the dynein occupying the trailing position on dynactin determines the velocity of two-motor assemblies, and that a trailing P3018S motor slows the entire complex regardless of the identity of the leading motor. We further demonstrate that the LIC of the leading dynein engages the trailing motor to increase its velocity by ∼130%, establishing a direct mechanism for leading-to-trailing motor stimulation within two-dynein DDB complexes. Together, these findings reveal how the P3018S mutation affects dynein function and uncover a positional and LIC-dependent mechanism that governs velocity control in multi-dynein assemblies.

## RESULTS

### The P3018S mutation resides in a regulatory AAA+ domain and corresponds to a naturally occurring variant in fungi lacking Lis1

Structural mapping places the P3018S substitution in dynein’s AAA4 module, adjacent to AAA3 (**Fig. 1b,c**). Although AAA1 is the principal ATPase that drives dynein stepping^46–52^, AAA3 and AAA4 play important regulatory roles in controlling conformational transitions within the AAA+ ring^47,53–56^. Cryo-EM studies further show that one β-propeller of the Lis1 homodimer binds directly to AAA4, while the second contacts the base of the stalk (**Fig. 1b**)^43,45^, positioning Lis1 to stabilize dynein in an open, active configuration^23,25^.

Because P3018 lies within the Lis1-interacting region of AAA4^23–31^, we considered whether this mutation disrupts Lis1-mediated activation. Unexpectedly, sequence alignments revealed that several *Schizosaccharomyces* species (*S. pombe*, *S. octosporus*, and *S. cryophilus*) naturally encode a serine at the position corresponding to P3018 (**Fig. 1d**). Notably, these same species lack a Lis1 ortholog^57^. Given that their dyneins support essential processes such as nuclear positioning and cell division^58–60^, their motors must perform without Lis1-dependent relief of autoinhibition.

These observations suggested that human dynein carrying the P3018S mutation may adopt an open conformation more readily than WT and thus display reduced dependence on Lis1 for activation.

### DDB-P3018S retains the ability to assume the phi conformation and respond to Lis1 but moves more slowly and generates less force

To test our hypothesis that P3018S destabilizes the autoinhibited state, we expressed recombinant WT dynein and the P3018S mutant in insect cells^19^, human dynactin in HEK293 suspension cells^61^, and human BicD2 in *E. coli*^61^ (**Supp. Fig. 1**). When assessed with single-molecule total internal fluorescence reflection (TIRF) microscopy, both WT and mutant motors formed processive DDB complexes that moved along surface-anchored microtubules when mixed at a 1:1:1 molar ratio^61^ (dynein:dynactin:BicD2) (**Fig. 2**).

**Figure 2.**
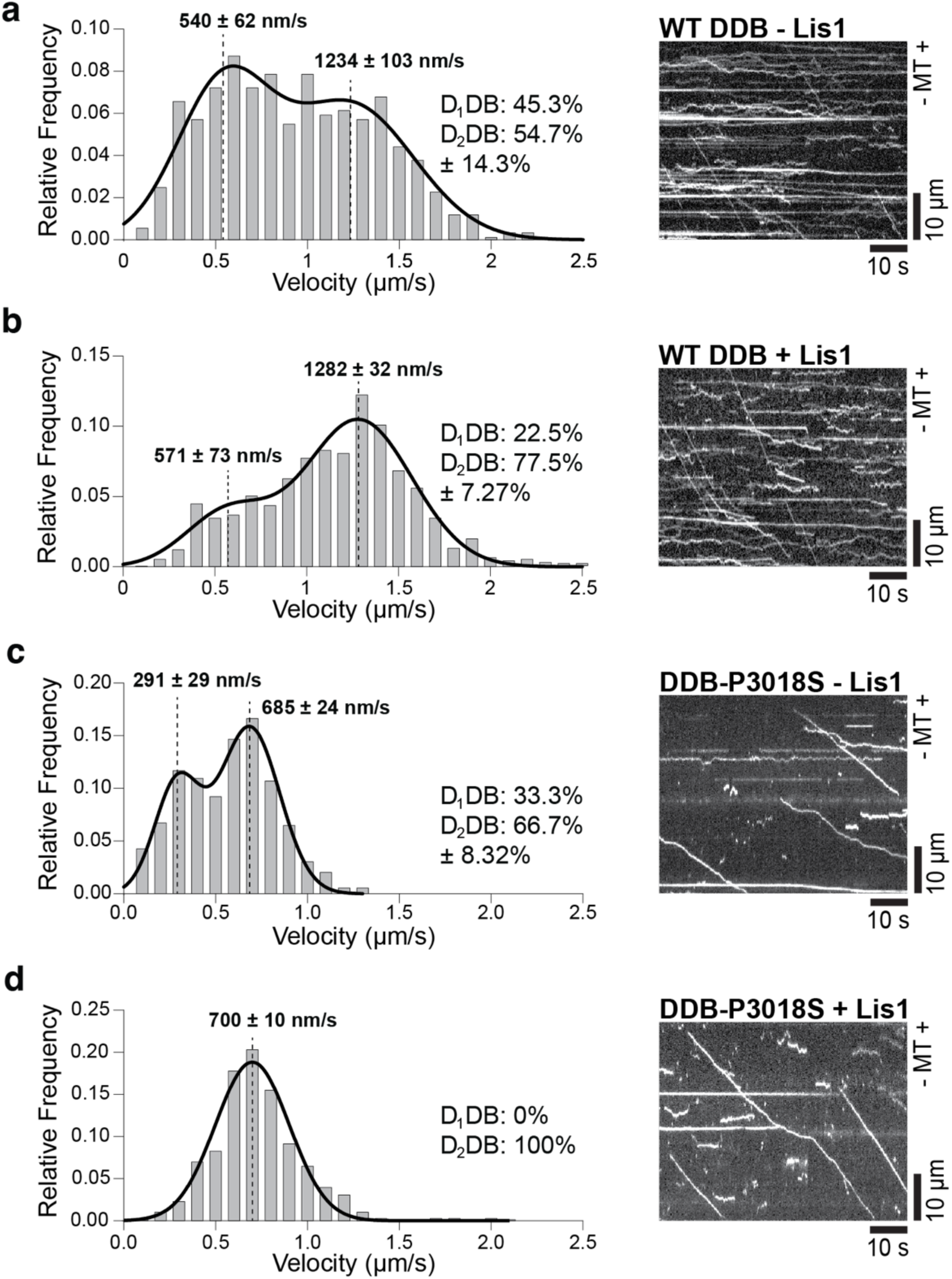
Effects of Lis1 and the P3018S mutation on DDB motility velocities. **(a-d)** Velocity distributions of processive, minus-end-directed motility events of dynein-dynactin-BicD2 (DDB) complexes under the indicated conditions. **(a)** Histogram of velocities measured for individual wild-type DDB complexes moving along microtubules in the absence of Lis1. The accompanying kymograph shows representative motility of Alexa Fluor 488–labeled wild-type dynein. Fitting the histogram with a two-component Gaussian mixture model (dashed lines) yielded the indicated mean velocities (± SEM) and the fractions of slow and fast motility events (± SEM) (*n* = 465). **(b)** Same as in (a), but for wild-type DDB complexes in the presence of Lis1 (*n* = 888). **(c)** Histogram of velocities measured for individual P3018S mutant DDB complexes (DDB-P3018S) moving along microtubules in the absence of Lis1. The accompanying kymograph shows representative motility of TMR-labeled P3018S dynein (*n* = 404). **(d)** Same as in (c), but in the presence of Lis1 (*n* = 788). In this condition, the velocity distribution was best described by a single Gaussian function.

For WT DDB, the velocity distribution was broad and described by two Gaussian components corresponding to single-dynein (D_1_DB) and two-dynein (D_2_DB) assemblies, centered near ∼600 and ∼1,200 nm/s, respectively, contributing ∼45% and ∼55% (**Fig. 2a**, left). Adding Lis1 increased the proportion of fast-moving events to ∼80% (**Fig. 2b**), consistent with Lis1 promoting formation of D_2_DB complexes^42,43^.

DDB-P3018S displayed markedly reduced velocities. The two Gaussian components were centered at ∼300 and ∼700 nm/s, and the higher-velocity component contributed ∼70% of events (70 ± 8% for DDB-P3018S vs. 55 ± 14% for WT; not significant, p ≈ 0.35) (**Fig. 2c**). Importantly, the mutant retained strong responsiveness to Lis1, with the fast-moving population approaching ∼100% upon Lis1 addition (**Fig. 2d**). The increased contribution of the higher-velocity component in the absence of Lis1 was qualitatively consistent with reduced entry into the autoinhibited phi conformation.

Negative-stain electron microscopy confirmed that P3018S dynein is capable of adopting both open and phi conformations (**Fig. 3a**). Although proportions varied between preparations, the presence of phi-state particles demonstrates that the mutant is not locked in an open conformation.

**Figure 3.**
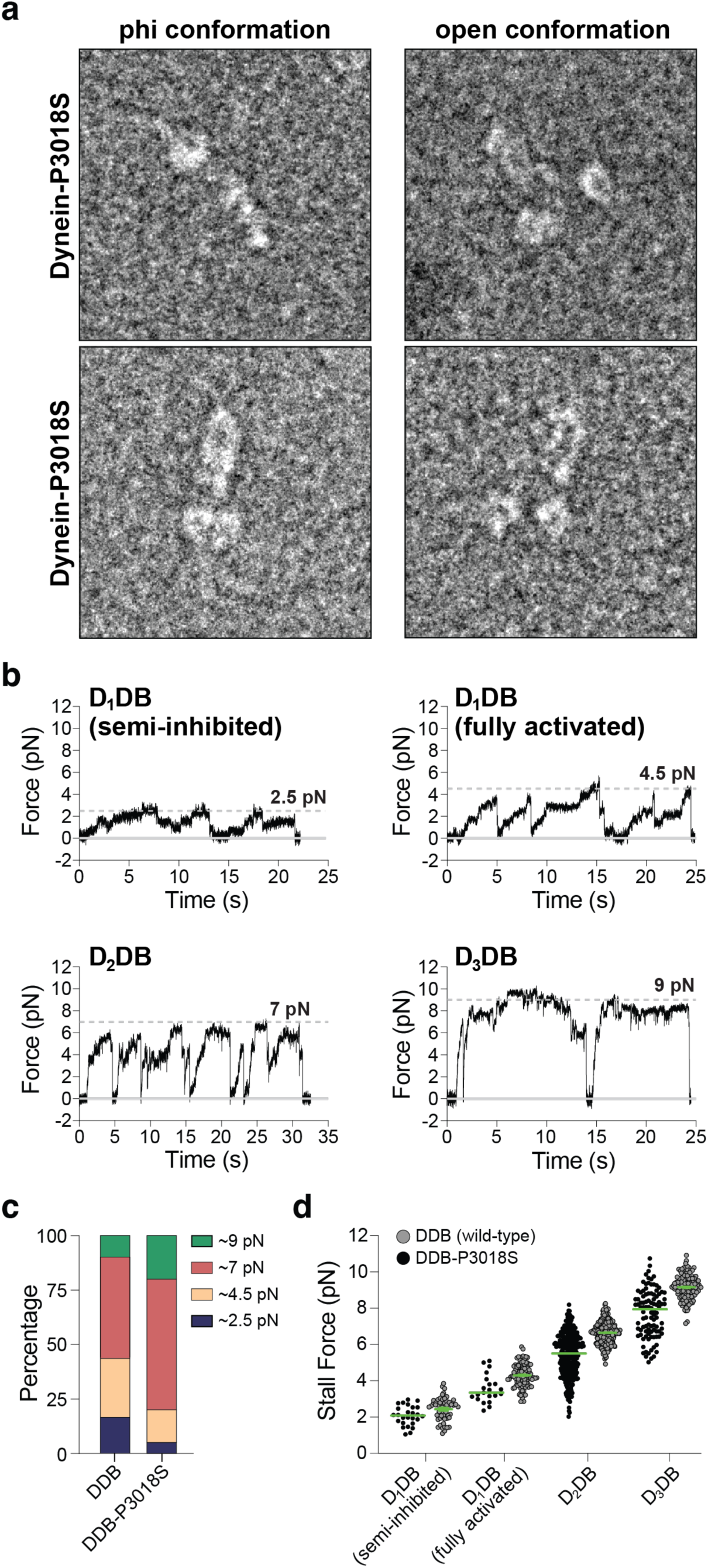
Conformational state and force generation of P3018S dynein–dynactin–BicD2 complexes. **(a)** Negative-stain electron microscopy micrographs of P3018S mutant dynein showing particles in the φ (phi) and open conformations. **(b)** Representative optical-tweezers traces illustrating force generation by DDB-P3018S complexes containing one, two, or three dyneins (D_1_DB, D_2_DB, and D_3_DB). Examples are shown for semi-inhibited and fully activated D_1_DB-P3018S, as well as for D_2_DB-P3018S and D_3_DB-P3018S. **(c)** Distribution of stall forces for wild-type and P3018S mutant dynein-dynactin-BicD complexes. Stall forces cluster at ∼2.5, ∼4.5, ∼7 and ∼9 pN and are shown for semi-inhibited D_1_DB, fully activated D_1_DB, D_2_DB, and D_3_DB assemblies. Percentages indicate the fraction of beads exhibiting each stall-force population. For wild-type DDB: *n* = 41 (17%, 27%, 46%, and 10%) (reproduced from ref^61^). For DDB-P3018S: *n* = 20 (5%, 15%, 60%, and 20%). **(d)** Stall forces measured for wild-type and P3018S mutant dynein–dynactin–BicD2 complexes containing one, two, or three dyneins (D_1_DB, D_2_DB, and D_3_DB). Data are shown as median values with 95% confidence intervals (CI). Semi-inhibited D_1_DB: *n* = 59, 2.6 [2.0, 2.8] pN; fully activated D_1_DB: *n* = 100, 4.2 [3.8, 4.8] pN; D_2_DB: *n* = 272, 6.7 [6.2, 7.1] pN; D_3_DB: *n* = 116, 9.2 [8.7, 9.6] pN. Semi-inhibited D_1_DB-P3018S: *n* = 26, 2.1 [1.8, 2.5] pN (Welch’s two-sample t-test vs. wild type based on mean and SD, *p* = 0.0085); fully activated D_1_DB-P3018S: *n* = 21, 3.3 [3.1, 3.8] pN (*p* = 3 × 10⁻^4^); D_2_DB-P3018S: *n* = 320, 5.5 [5.3, 5.8] pN (p < 10⁻^30^); D_3_DB-P3018S: *n* = 100, 7.9 [7.2, 8.2] pN (p < 10⁻^15^).

To examine force generation, we performed single-molecule optical trapping. As in WT, DDB-P3018S displayed stalling events corresponding to one-, two-, and three-dynein assemblies^61^ (**Fig. 3b**). However, stall forces were consistently reduced for the mutant due to an increased detachment rate before a stalling plateau was reached. The fraction of ∼2.5 pN stalling events—indicative of entry into a phi-like state^61^—also tended to be lower for DDB-P3018S than for WT (5% vs. 17% for WT, p < 0.09; **Fig. 3c**), consistent with a modest reduction in the frequency of autoinhibition.

Thus, the P3018S mutant retains phi-state formation, Lis1 responsiveness, and load-dependent recruitment of additional motors but exhibits slower velocity and reduced force output. The reduced frequency of phi-like stalls indicates a shift in the equilibrium away from autoinhibition rather than loss of the inhibited state.

### The trailing dynein motor determines the velocity of two-dynein DDB complexes

Because we currently cannot generate full-length heterodimeric dynein containing one WT and one P3018S heavy chain, we reconstituted mixed-motor DDB assemblies comprising one homodimeric WT dynein and one homodimeric P3018S dynein. In a heterozygous patient, dynein populations are expected to include WT homodimers, mutant homodimers, and WT–mutant heterodimers, which can combine to form DDB complexes with different two-motor configurations. Our mixed assemblies therefore model a subset of physiologically relevant WT–mutant pairings in which the two motors operate together on the same DDB complex.

To determine how coupling a fast WT motor to a slower mutant affects ensemble motility, we differently labeled WT dynein (Alexa Fluor 488) and P3018S mutant dynein (TMR) and performed dual-color single-molecule TIRF microscopy (**Fig. 4a-d**). Co-localization of both fluorophores confirmed that the analyzed complexes contained two dyneins. The velocity distribution of mixed assemblies was well fit by two Gaussian components centered at ∼550 and ∼1,000 nm/s. Because Lis1 primarily promotes recruitment of a second dynein and the analyzed complexes already contained two motors, Lis1 addition did not alter the relative fractions of slow and fast events (**Fig. 4a,b**). Thus, these two velocity classes represent intrinsic properties of the mixed WT–mutant assemblies rather than reflecting differences in motor copy number.

**Figure 4.**
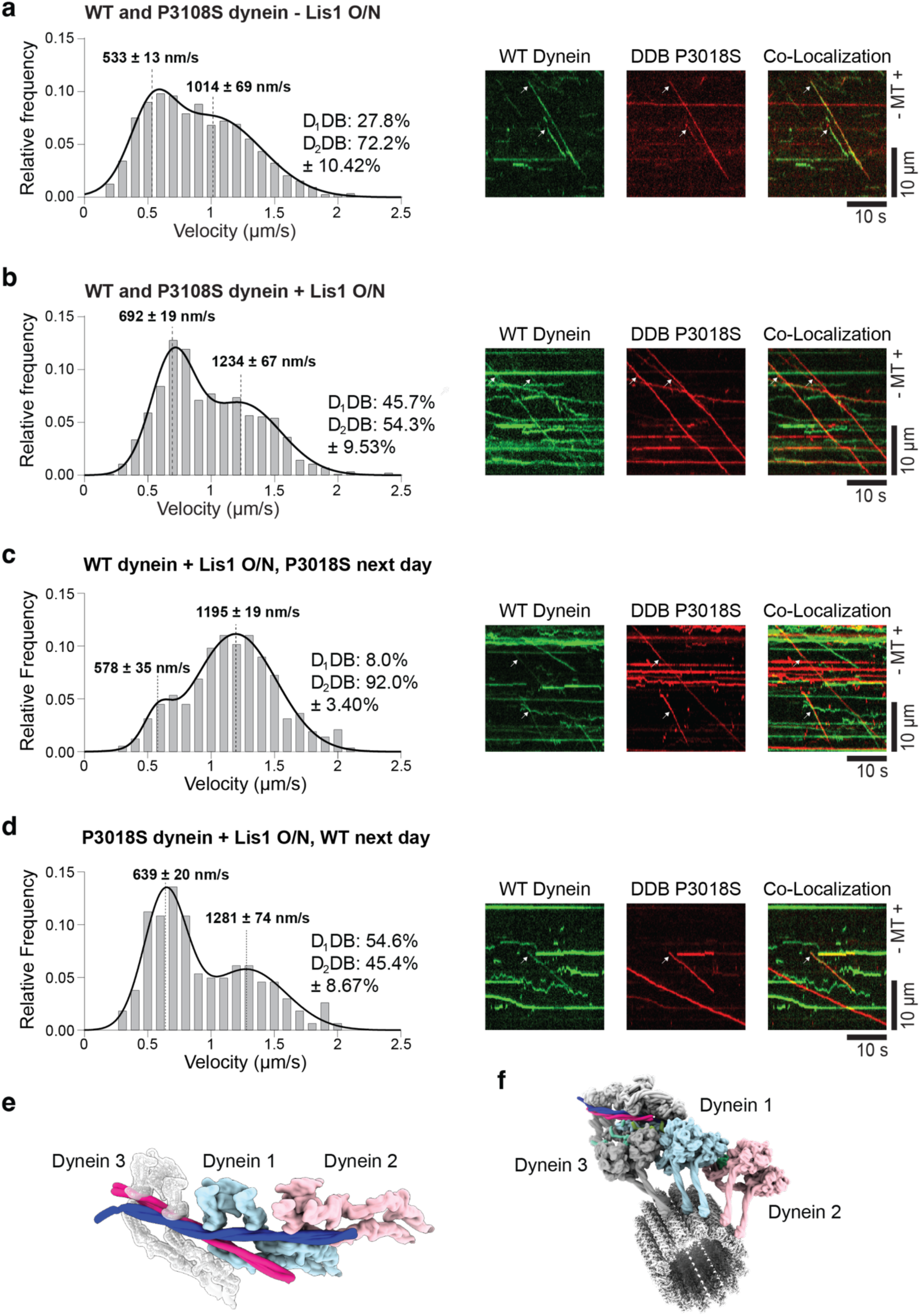
Structural basis and motility of mixed wild-type/P3018S dynein DDB complexes. (**a-d**) Velocity distributions of processive, minus end–directed motility events for DDB complexes containing one Alexa Fluor 488–labeled wild-type dynein and one TMR-labeled P3018S mutant dynein moving together along microtubules, in the absence or presence of Lis1. Representative kymographs show co-localized motility of both dyneins. Velocity histograms were fit with two-component Gaussian mixture models (dashed curves) to obtain mean velocities (± SEM) and the fractions of slow and fast motility events (± SEM) under the indicated incubation conditions. (**a**) DDB complexes assembled overnight (ON) with wild-type dynein and P3018S mutant dynein in the absence of Lis1 (*n* = 990). (**b**) Same as in (a), but with Lis1 present during assembly (*N* = 828). (**c**) DDB complexes assembled ON with wild-type dynein and Lis1, followed by addition of P3018S dynein on the following day (*n* = 582). (**d**) DDB complexes assembled ON with P3018S dynein and Lis1, followed by addition of wild-type dynein on the following day (*n* = 255). (**e**) Cryo-EM structure of the dynein–dynactin–BICDR1 (DDR) complex (PDB: 7Z8F) showing the staggered binding of the tails of two dyneins (Dynein 1 and Dynein 2) to the cargo adaptors and dynactin. A semi-transparent rendering with gray helices indicates the predicted binding site for a third dynein. To visualize the predicted interaction with BicD2 rather than BICDR1 and to improve clarity, an AlphaFold3 model of BicD2(18–240) was superimposed onto the BICDR1 position; BICDR1 and dynactin were subsequently hidden for visualization. (**f**) Model of the full DDR assembly including a third dynein (gray) and a microtubule.

In our previous WT-only or P3018S-only experiments, the slow and fast peaks corresponded to one- and two-dynein assemblies, respectively (**Fig. 2**). In contrast, all mixed assemblies contain two dyneins, indicating that the two discrete velocities must arise from positional differences. Structural studies show that the two dyneins occupy staggered positions on the dynactin shoulder: a trailing motor (Dynein 1, **Fig. 4e**) that binds first via the adaptor^32,62^, and a leading motor (Dynein 2, **Fig. 4e**) that binds second^33,36^; a third motor (Dynein 3, **Fig. 4e**) only associates under load^61^. If this staggered arrangement biases control toward the trailing motor, then the identity of the trailing dynein should disproportionately influence the velocity of the two-motor assembly. A trailing WT could maintain high velocity even when paired with a mutant in the leading position, whereas a trailing mutant could slow the entire complex.

To test this model, we took advantage of the sequential loading of dynein onto dynactin^33,62^. We first assembled DDB complexes overnight using WT dynein alone, conditions expected to populate dynactin with a WT trailing motor^62^ (Dynein 1, **Fig. 4e**) and, where present, a WT leading motor^33^ (Dynein 2, **Fig. 4e**). The next day, we added P3018S dynein for one hour. Because Dynein 1 is likely stably bound over hours whereas Dynein 2 exchanges more rapidly^33,62^, this protocol should preferentially position the mutant in the leading site while maintaining a WT trailing motor. Conversely, when the overnight incubation was performed with P3018S dynein and WT dynein was added the next day, the mutant should occupy the trailing position with the WT motor preferentially loading into the leading position.

This positional manipulation produced a striking shift in velocity distributions. When WT dynein was used for the overnight incubation (WT in the trailing position), the slow ∼600 nm/s peak represented only 8 ± 3% (mean ± SEM) of events (**Fig. 4c**). When the overnight incubation was done with P3018S dynein (mutant in the trailing position), the slow peak increased to 55 ± 9% (*p* < 10^−5^) (**Fig. 4d**). These data strongly support the model that the trailing motor dictates the velocity of two-dynein DDB complexes. Moreover, complexes assembled with a mutant in the trailing position move at the mutant two-dynein velocity regardless of whether the leading motor is WT or mutant. Consistent with this, two-dynein P3018S-only complexes move at 700 ± 10 nm/s, comparable to the 639 ± 20 nm/s velocity of mixed assemblies containing one WT and one mutant dynein.

### The leading dynein’s LIC accelerates the trailing motor by ∼130%

Recent cryoEM studies suggests that the LIC of the leading motor engages AAA2/AAA3 of the trailing motor^63^ (**Fig. 5a**), providing a structural rationale for how the leading motor could enhance trailing-motor activity. To test whether this interaction underlies the ∼130% velocity increase associated with the trailing dynein, we took advantage of DDB complexes containing a full-length dynein paired with a dynein tail fragment that retains all associated chains, including the LIC, but lacks the motor domain^35^. Overnight incubation with WT dynein and the LIC-containing tail fragment (the reverse order yielded too few moving events for analysis) produced DDB assemblies in which a single motor domain is paired with a tail-only “leading” unit. As previously shown^35^, these hybrid complexes move at the velocity of a two-dynein DDB assembly (**Fig. 5b–d**).

**Figure 5.**
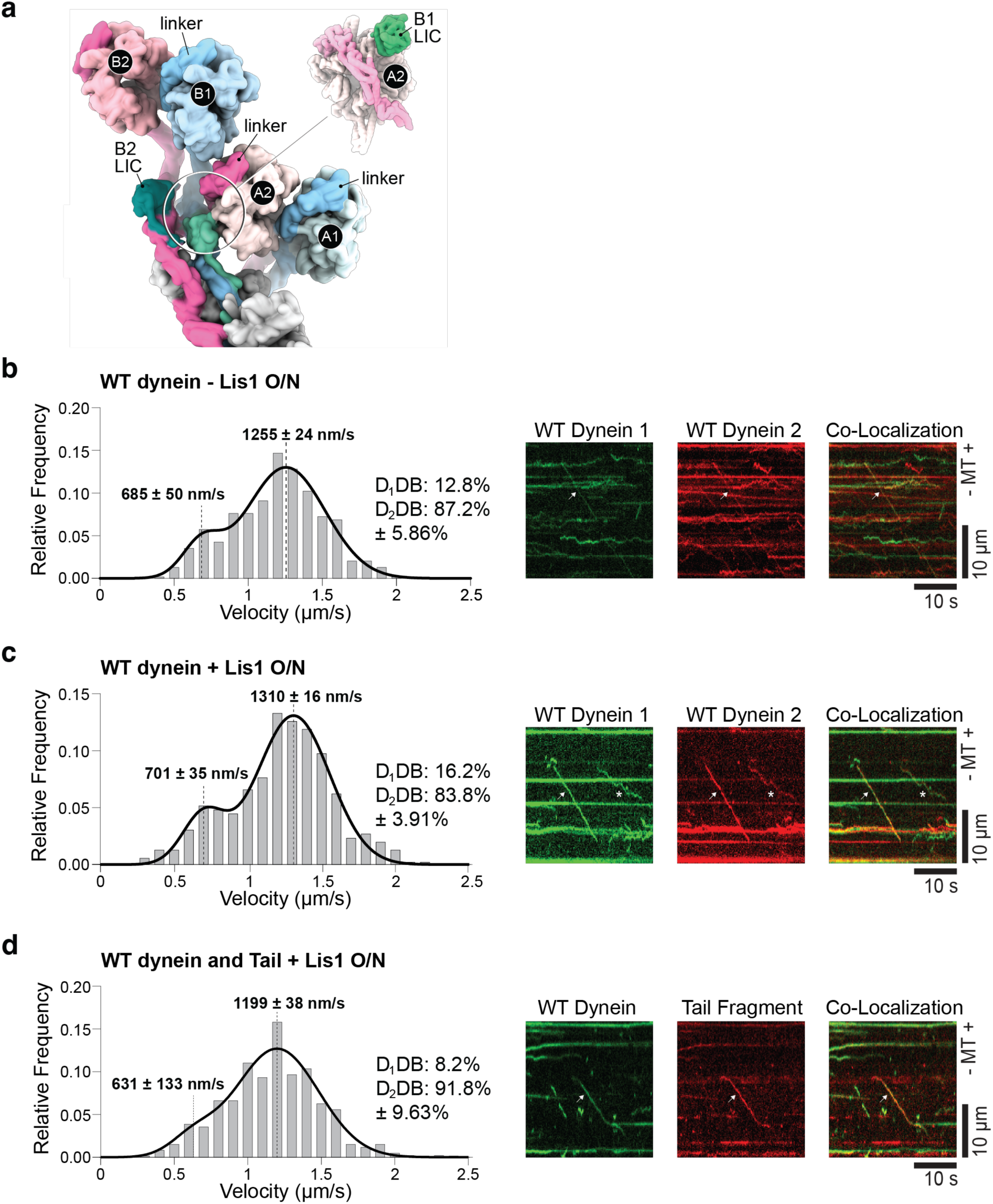
LIC-mediated interactions between leading and lagging dyneins enhance DDB motility. (**a**) Top-view surface representation of the dynein–dynactin–JIP3 (DDJ) complex containing two dyneins, highlighting the molecular interface between the light intermediate chain (LIC) of the right-hand heavy chain of the leading dynein (B2) and the left-hand motor domain of the lagging dynein (A2) (PDB: 7Z8F). The inset shows a surface view of the interacting regions of dynein A2 and the LIC of dynein B2. (**b**-**d**) Velocity distributions of processive, minus end–directed motility events for DDB complexes assembled overnight (ON) under the indicated conditions. Representative kymographs show co-localized motility of both dyneins or motility of DDB complexes containing a single dynein and a dynein tail fragment. Velocity histograms were fit with two-component Gaussian mixture models (dashed curves) to obtain mean velocities (± SEM) and the fractions of slow and fast motility events (± SEM). (**b**) DDB complexes containing one Alexa Fluor 488–labeled wild-type dynein and one TMR-labeled wild-type dynein in the absence of Lis1 (*n* = 269). (**c**) Same as in (b), but in the presence of Lis1 (*n* = 283). (**d**) DDB complexes containing one Alexa Fluor 488–labeled wild-type dynein and a TMR-labeled dynein tail domain (*n* = 294).

Notably, even when two dyneins were present on the complex—as confirmed by two-color co-localization—the velocity distribution remained biphasic (**Fig. 5b**). Lis1 addition did not alter the relative proportions of the two Gaussian components (**Fig. 5c**), consistent with our earlier conclusion that the two velocities arise from intrinsic motor-state dynamics rather than from differences in motor copy number. These data suggest that stimulation of the trailing dynein by the LIC-containing leading unit is dynamic: the complex spends part of its time in a high-velocity, LIC-stimulated state and part in a single-dynein–like state near ∼600 nm/s.

Additional evidence comes from the behavior of three-dynein DDB complexes^61^. Rather than increasing velocity via load sharing, recruitment of a third dynein—which occupies the most trailing position on the dynactin shoulder (**Fig. 4e,f**)—slows the complex to speeds even lower than those of a single-dynein DDB assembly^61^. This further supports the conclusion that the trailing dynein in the staggered arrangement is the dominant determinant of ensemble velocity.

## DISCUSSION

Cytoplasmic dynein must integrate regulatory cues, mechanical load, and multi-motor coordination to support long-range intracellular transport. Here we define how a patient-associated mutation in the AAA+ ring perturbs these regulatory layers and uncover a positional mechanism by which dynein motors influence one another within dynein–dynactin–adaptor assemblies.

P3018S resides in AAA4 (**Fig. 1b**), a regulatory module positioned adjacent to AAA3 and forming part of the Lis1-interacting surface^45,64^. Given that several *Schizosaccharomyces* species lacking Lis1^57^ naturally encode a serine at this position, we initially hypothesized that P3018S might bias human dynein toward an activated state. Instead, we find that P3018S slows dynein stepping and reduces force output while preserving key regulatory features: the mutant motor maintains the capacity to enter the autoinhibited phi conformation, responds robustly to Lis1, and assembles into higher-order force-generating states. Notably, however, quantitative analysis shows that the P3018S mutant exhibits a reduced probability of adopting a phi-like conformation, indicating that the mutation perturbs the equilibrium between inhibited and active conformations without eliminating access to either. Taken together, these results show that P3018S does not abolish dynein’s ability to undergo Lis1-regulated transitions but instead alters the likelihood that these transitions occur.

A central discovery of this study is that ensemble velocity in two-dynein DDB complexes depends on which motor occupies the trailing position on dynactin. Mixed wild-type–mutant reconstitutions revealed two velocity populations despite every complex containing two dyneins, demonstrating that differences in motor number alone cannot explain the observed behavior. Instead, the identity of the trailing dynein dictates the speed of the entire complex: a trailing wild-type motor supports wild-type two-dynein velocities, whereas a trailing P3018S motor slows the complex to the mutant two-dynein velocity, independent of the leading motor. This positional asymmetry provides experimental support for models proposing that dynein’s staggered arrangement on dynactin generates intrinsic functional hierarchy among motors^61^.

Our data further reveal that the leading dynein actively stimulates the trailing dynein through a structural interaction mediated by the light-intermediate chain (LIC)^63^. Using tail-only constructs that retain all associated dynein subunits, including the LIC, but lack the motor domain, we found that LIC alone is sufficient to support the ∼130% increase in trailing-motor velocity normally observed in two-dynein assemblies. These experiments demonstrate that LIC-dependent contacts between the leading and trailing dyneins dynamically enhance trailing-motor activity, establishing a mechanism for leading-to-trailing motor stimulation. These results indicate that dyneins within the same complex do not operate as independent motors but instead regulate one another through defined structural interfaces, revealing intra-complex allosteric communication. This LIC-mediated regulatory step adds a layer to dynein activation distinct from adaptor-driven recruitment and Lis1-mediated relief of autoinhibition.

Recent work further supports a central regulatory role of the LIC within multi-motor dynein–dynactin assemblies. In our accompanying study^61^, we show that an auxiliary BicD2 adaptor binds the LIC of a third dynein, enabling tension-dependent recruitment of an additional motor to DDB complexes and thereby increasing ensemble force output. Together with the present finding that LIC-dependent contacts between the leading and trailing dyneins potentiate the activity of the trailing motor, these results support a directional role for the LIC in coordinating dynein ensembles. Notably, however, DDB complexes containing three dyneins move more slowly than two-dynein assemblies under load^61^, despite the clear stimulatory effect of the leading dynein’s LIC in two-dynein DDB. This non-monotonic relationship between motor number and velocity argues against a simple additive LIC-mediated activation mechanism and instead suggests that LIC-dependent stimulation may be restricted to the immediately trailing dynein and is not efficiently propagated to additional motors. Such a limitation may reflect geometric constraints imposed by the dynactin scaffold or redistribution of inter-motor tension in higher-order assemblies. Thus, the LIC appears to serve a dual, context-dependent function within dynein–dynactin complexes: the LIC of the leading motor transmits stimulatory input to the motor directly behind it, whereas the LIC of the terminal motor provides a docking interface for auxiliary adaptors that nucleate higher-order assemblies. Consistent with this model, structural and biochemical studies show that adaptor CC1-box motifs engage conserved LIC helices^63,65^ and that LIC-adaptor affinity is tunable through defined interaction surfaces and regulated by cargo context and phosphorylation^22,66^.

Together, these results reveal that dynein–dynactin assemblies employ positional control and inter-motor communication to tune their motile properties. The trailing motor functions as the primary determinant of ensemble speed, while the leading motor contributes regulatory input through LIC-dependent stimulation. These principles help explain why dynein complexes with identical motor composition can display heterogeneous velocities and offer a framework for understanding how cargo adaptors, cofactors, and mechanical load shape dynein-driven transport. Our findings refine prevailing models by demonstrating that the two dyneins bound to dynactin make unequal functional contributions, with distinct roles in force generation and motility control.

For disease biology, our findings provide mechanistic insight into how mutations in AAA4 may impair dynein-driven trafficking. Although P3018S preserves dynein’s ability to form the activated DDB state, its reduced stepping velocity and force generation, combined with its dominant influence when occupying the trailing position, suggest that even partial loss-of-function variants may disproportionately impact ensemble behavior in cells. Given the prevalence of heterozygous DYNC1H1 mutations in neurodevelopmental disorders^9–11,13,15–17,67–74^, positional dominance within multi-motor assemblies may represent a general mechanism by which subtle biochemical defects translate into cellular dysfunction.

In summary, this work elucidates how a disease-associated AAA4 mutation influences dynein’s mechanochemistry and reveals that dynein–dynactin motility emerges from the integrated activities of motors in distinct positional states. The identification of LIC-dependent leading-to-trailing stimulation provides a mechanistic basis for cooperative activation within multi-motor assemblies and introduces a positional paradigm for dynein regulation with implications for neuronal transport, motor coordination, and dyneinopathies.

## Materials and Methods

### Plasmid and Construct Generation

Plasmids encoding human dynein complexes were generously provided by Dr. Andrew P. Carter (MRC Laboratory of Molecular Biology, Cambridge, UK). Plasmids encoding Lis1 were gifts from Dr. Samara Reck-Peterson (University of California, San Diego, CA).

BicD2 constructs were generated by subcloning mouse *BicD2*(25–400) from an existing plasmid (Addgene #64205)^75^ into an *E. coli* expression backbone (pSNAP-tag(T7)-2; New England Biolabs, #N9181S). The SNAP tag was subsequently replaced with either a StrepII tag or an Avi–StrepII tag using Q5 site-directed mutagenesis (New England Biolabs, #E0554S). Plasmids encoding TEV protease (Addgene #171782)^76^ and BirA (Addgene #20857)^77^ were obtained from Addgene.

The P3018S mutation was introduced into the human dynein heavy chain (DHC) using a previously published strategy^78^. Briefly, the mutation was generated in the DHC-containing plasmid pbiG1a:6His-ZZ-SNAPf-DHC1 by Gibson assembly. The resulting PmeI-digested gene expression cassette was co-assembled with the PmeI-digested polycistronic cassette from pbiG1b:IC2/LIC2/Tctex1/Robl1/LC8 into PmeI-digested pbiG2ab using biGBac cloning techniques^79^. All pbiG vectors were kindly provided by Dr. Steven M. Markus (Colorado State University, USA).

All plasmids generated in this study were validated by Sanger sequencing (Genomics Core Facility, Albert Einstein College of Medicine, Bronx, NY) or by full-plasmid sequencing (Azenta).

### HEK Suspension Cell Culture

One milliliter of HEK293T/17 SF cells (ATCC, ACS-4500) expressing C-terminally tagged DCTN4 (p62-HaloTag-3×FLAG-P2A-mCherry) was recovered according to the manufacturer’s instructions and diluted into 10 mL of complete medium. The complete medium consisted of 9.8 mL BalanCD HEK293 (Fujifilm Irvine Scientific, #91165-1L), 0.1 mL of 100× GlutaMAX™ supplement (Gibco, #35050061), 0.1 mL of 100× ITS (Insulin-Transferrin-Selenium; Corning, #25-800-CR), and 20 µL of anti-clumping agent (Gibco, #0010057DG).

Cultures were maintained at 37 °C with 5% CO₂ under constant agitation at 125 rpm in a CO₂ incubator shaker (New Brunswick S41i, Eppendorf) and kept at a cell density below 2 × 10⁶ cells/mL.

### Dynactin Purification

HEK293T/17 SF cells expressing p62 tagged with HaloTag-3×FLAG were cultured in 800 mL of medium at a density of 2 × 10⁶ cells/mL and harvested by centrifugation at 600 × g for 10 minutes at 4 °C. The resulting cell pellet (∼10 mL) was resuspended in 10 mL of 2× dynactin lysis buffer (DLB) containing 60 mM HEPES (pH 7.2), 200 mM NaCl, 4 mM MgCl₂, 2 mM EGTA, 20% glycerol, 0.2 mM ATP, 2 mM DTT, 0.4% (v/v) Triton X-100, and two EDTA-free protease inhibitor cocktail tablets (Roche, #11836170001). The suspension was nutated at 4 °C for 15 minutes. The lysate was clarified by centrifugation at 80,000 rpm (260,000 × g, k-factor = 28) for 10 minutes using a TLA-110 rotor in a Beckman Optima TLX tabletop ultracentrifuge.

Anti-FLAG® M2 affinity gel (1 mL; Sigma, #A2220) was washed with 5 mL of 1× DLB. The cleared lysate was added to the resin and nutated overnight at 4 °C. The resin was subsequently washed with 40 mL of wash buffer containing 30 mM HEPES (pH 7.2), 250 mM KCl, 2 mM MgCl₂, 1 mM EGTA, 10% glycerol, 0.1 mM ATP, 1 mM DTT, 0.2% (w/v) Pluronic F-127, and 0.5 mM Pefabloc.

The resin was then transferred to a 2-mL tube and incubated with 200 µL of wash buffer supplemented with 100 µL of 5 mg/mL FLAG peptide. The mixture was nutated at 4 °C for 30 minutes. The supernatant containing purified dynactin was collected and concentrated using an Amicon Ultra 0.5-mL centrifugal filter unit (100-kDa MWCO) at 5,000 × g and 4 °C. Dynactin concentration was determined using a Bradford assay (Thermo Fisher Scientific, #23200), and subunit composition was confirmed by mass spectrometry.

### Sf9 Expression of Cytoplasmic Dynein-1 and Lis1

The full cytoplasmic dynein-1 complex, the P3018S mutant, and Lis1 were expressed in Sf9 cells following established protocols^19^. Briefly, bacmids were generated by transforming plasmids encoding the desired constructs into MAX Efficiency™ DH10Bac competent cells (Gibco, #10361012). Recombinant bacmid insertion was verified by blue-white screening and confirmed by PCR.

One milliliter of Sf9 cells cultured in Sf-900™ II SFM (Thermo Fisher Scientific, #11496015) was recovered and expanded in 10 mL of Sf-900™ II SFM medium (Thermo Fisher Scientific, #10902104) at 27 °C with shaking at 135 rpm. Cultures were maintained at a density below 2 × 10⁶ cells/mL. For transfection, 2 mL of a culture at 0.5 × 10⁶ cells/mL was seeded into a 6-well plate. A mixture containing 2 µg of bacmid DNA and 200 µL of medium was prepared, and 6 µL of FuGENE® HD transfection reagent (Promega, #E2311) was added. After incubation for 15 minutes at room temperature, the mixture was added dropwise to the cells.

The cultures were then incubated at 27 °C without shaking for 4 days to generate P1 virus. The P1 virus was collected, transferred to a 15-mL tube, and stored at 4 °C in the dark. To produce P2 virus, 0.5 mL of P1 virus was added to 50 mL of Sf9 cells at a density of 1.5 × 10⁶ cells/mL and incubated at 27 °C with shaking at 135 rpm for 3 days. The P2 virus was harvested by centrifugation at 2,000 × g for 10 minutes at 4 °C and stored at 4 °C in the dark.

For protein expression, 5 mL of P2 virus was added to 500 mL of Sf9 cells at a density of 2 × 10⁶ cells/mL. Cultures were incubated at 27 °C with shaking at 135 rpm for 60–72 hours. Cells were harvested by centrifugation at 2,000 × g for 15 minutes at 4 °C. The resulting pellet was washed with 40 mL of cold PBS and centrifuged again at 2,000 × g for 10 minutes at 4 °C. The supernatant was discarded, and the pellet was flash-frozen in liquid nitrogen and stored at −80 °C until further use.

### Cytoplasmic Dynein-1 and Lis1 Purification

The cell pellet (∼5 mL) was resuspended in 5 mL of 2× lysis buffer containing 60 mM HEPES (pH 7.2), 300 mM KCl, 4 mM MgCl₂, 2 mM EGTA, 20% (v/v) glycerol, 0.4 mM ATP, 4 mM DTT, 0.2% (w/v) Pluronic F-127, 100 µL of 2.5 U/µL DNase I, and two EDTA-free protease inhibitor cocktail tablets. The resuspended pellet was dounced with 30 strokes on ice. The lysate was cleared by centrifugation as described for dynactin purification.

Four milliliters of IgG Sepharose 6 Fast Flow affinity resin (Cytiva, #17096901) was pre-washed with 6 mL of lysis buffer. The cleared lysate was added to the resin and nutated at 4 °C for 4 hours. The resin was then washed with 50 mL of wash buffer containing 50 mM HEPES (pH 7.2), 150 mM KCl, 2 mM MgCl₂, 1 mM EGTA, 10% (v/v) glycerol, 0.1 mM ATP, 1 mM DTT, 0.1% (w/v) Pluronic F-127, and 1 mM PMSF.

To label the SNAP tag at the N-terminus of the dynein heavy chain, 15 µL of 1 mM SNAP-tag dye was added directly to the resin, and the mixture was nutated at 4 °C overnight. The resin was then washed with 50 mL of TEV-release buffer containing 50 mM HEPES (pH 7.2), 150 mM KCl, 2 mM MgCl₂, 1 mM EGTA, 10% (v/v) glycerol, 0.1 mM ATP, 1 mM DTT, and 0.1% (w/v) Pluronic F-127.

The resin was subsequently transferred to a 5-mL tube and resuspended in TEV-release buffer to a final volume of 5 mL. Fifty microliters of 200 µM purified MBP-superTEV protease was added, and the mixture was nutated at 4 °C overnight in the dark. One milliliter of amylose resin (New England Biolabs, #E8021S) was used to remove the MBP-superTEV protease. The supernatant was concentrated following the same protocol used for dynactin purification.

The purity of the cytoplasmic dynein-1 complex and Lis1 was assessed by 4–12% Bis-Tris polyacrylamide gel electrophoresis, and protein concentration was determined using a Bradford assay. WT dynein was labeled with SNAP-surface Alexa Fluor 488 (New England Biolabs, #S9129S), and mutant dynein was labeled with SNAP-cell TMR Star488 (New England Biolabs, #S9105S). Dynein tail was labeled with either SNAP-cell TMR Star or SNAP-surface Alexa Fluor 647 (New England Biolabs, #S9136S).

### *E. coli*-Based Expression

BicD2 constructs were expressed in *E. coli*. Each plasmid was transformed into BL21-CodonPlus(DE3)-RIPL competent cells (Agilent Technologies, #230280). A single colony was picked and inoculated into 1 mL of terrific broth (TB) containing 50 µg/mL carbenicillin and 50 µg/mL chloramphenicol. The 1-mL starter culture was shaken overnight at 37 °C and subsequently used to inoculate 400 mL of TB supplemented with 2 µg/mL carbenicillin and 2 µg/mL chloramphenicol.

The culture was shaken at 37 °C for 5 hours and then cooled to 16 °C for 1 hour. Protein expression was induced with 0.1 mM IPTG and continued overnight at 16 °C. Cells were harvested by centrifugation at 3,000 × g for 10 minutes at 4 °C, and the supernatant was discarded. The cell pellet was resuspended in 5 mL of B-PER™ Complete Bacterial Protein Extraction Reagent (Thermo Fisher Scientific, #89821) supplemented with 2 mM MgCl₂, 1 mM EGTA, 1 mM DTT, 0.1 mM ATP, and 2 mM PMSF. The resuspension was flash-frozen in liquid nitrogen and stored at −80 °C.

### Purification of *E. coli*-Expressed Constructs

To purify *E. coli*–expressed proteins, frozen cell pellets were thawed at 37 °C and nutated at room temperature for 20 minutes to lyse the cells. The lysate was cleared as described for dynactin purification. The resulting supernatant was passed through 500 µL of Ni-NTA cOmplete™ His-Tag purification resin (Millipore Sigma, #5893682001) for His-tagged proteins or through Strep-Tactin® 4Flow® high-capacity resin (IBA Lifesciences GmbH, #2-1250-010) for StrepII-tagged proteins.

The resin was washed with 10 mL of wash buffer containing 50 mM HEPES (pH 7.2), 300 mM KCl, 2 mM MgCl₂, 1 mM EGTA, 1 mM DTT, 1 mM PMSF, 0.1 mM ATP, 0.1% (w/v) Pluronic F-127, and 10% glycerol. Proteins were eluted with elution buffer containing 50 mM HEPES (pH 7.2), 150 mM KCl, 2 mM MgCl₂, 1 mM EGTA, 1 mM DTT, 1 mM PMSF, 0.1 mM ATP, 0.1% (w/v) Pluronic F-127, and 10% glycerol, supplemented with either 150 mM imidazole for His-tagged proteins or 5 mM desthiobiotin for StrepII-tagged proteins.

Eluted proteins were either flash-frozen and stored at −80 °C or concentrated using an Amicon Ultra 0.5-mL centrifugal filter unit (30-kDa MWCO; Sigma, #UFC503024) prior to flash freezing. Protein purity and concentration were assessed using 4–12% Bis-Tris polyacrylamide gel electrophoresis.

For biotinylation of BicD2, 50 µL of 10 µM BicD2 was mixed with 1 µL of 100 mM ATP, 2 µL of 50 µM purified BirA, and 1 µL of 10 mM biotin and incubated at 30 °C for 2 hours. BirA was subsequently removed using Ni-NTA resin, and the solution was buffer-exchanged into 30 mM HEPES (pH 7.2), 50 mM KCl, 2 mM MgCl₂, 1 mM EGTA, 1 mM DTT, 0.1% (w/v) Pluronic F-127, and 10% glycerol to remove free biotin.

### Optical Tweezers Assay

#### Polystyrene Beads

Streptavidin beads (0.55 µm) were purchased from Spherotech (#SVP-05-10).

#### Microtubule Polymerization

Two microliters of 10 mg/mL tubulin (Cytoskeleton, #T240-B) were mixed with 2 µL of 1 mg/mL biotinylated tubulin (Cytoskeleton, #T333P-A) and 1 µL of 10 mM GTP. The mixture was incubated at 37 °C for 20 minutes. Following incubation, 0.5 µL of 0.2 mM paclitaxel in DMSO was added, and incubation was continued for an additional 20 minutes.

The solution was carefully layered onto a 100-µL glycerol cushion containing 80 mM PIPES (pH 6.8), 2 mM MgCl₂, 1 mM EGTA, 60% (v/v) glycerol, 1 mM DTT, and 10 µM paclitaxel in a 230-µL TLA100 tube (Beckman Coulter, #343775) and centrifuged at 80,000 rpm (250,000 × g, k-factor = 10) for 5 minutes at room temperature. The supernatant was gently removed, and the pellet was resuspended in 11 µL of BRB80G10 buffer containing 80 mM PIPES (pH 6.8), 2 mM MgCl₂, 1 mM EGTA, 10% (v/v) glycerol, 1 mM DTT, and 10 µM paclitaxel. The microtubule solution was stored at room temperature in the dark until further use.

#### Flow Chamber Preparation

Flow chambers were assembled using a glass slide (Fisher Scientific, #12-550-123) and an ethanol-cleaned coverslip (Zeiss, #474030-9000-000), separated by two thin strips of parafilm. Ten microliters of 0.5 mg/mL BSA-biotin (Thermo Scientific, #29130) were introduced into the chamber and incubated for 10 minutes.

The chamber was then washed with 2 × 20 µL of blocking buffer containing 80 mM PIPES (pH 6.8), 2 mM MgCl₂, 1 mM EGTA, 10 µM paclitaxel, 1% (w/v) Pluronic F-127, 2 mg/mL BSA, and 1 mg/mL α-casein, and incubated for 30 minutes to block the surface. Next, 10 µL of 0.25 mg/mL streptavidin (Promega, #Z7041) was introduced into the chamber and incubated for 10 minutes.

The chamber was washed with 2 × 20 µL of blocking buffer, followed by the addition of 10 µL of 0.02 mg/mL biotin-labeled microtubules in blocking buffer, which were incubated for 1 minute. After a final wash with 2 × 20 µL of blocking buffer, the chamber was stored in a humidity chamber until further use.

#### Sample Preparation

Dynein (dimer), dynactin, and BicD2 (dimer) were mixed at a 1:1:1 molar ratio to a final concentration of 200 nM each. The mixture was incubated at 4 °C overnight to allow complex formation.

One microliter of polystyrene trapping beads was mixed with 1 µL of motility buffer containing 60 mM HEPES (pH 7.2), 50 mM KCl, 2 mM MgCl₂, 1 mM EGTA, 10 µM paclitaxel, 0.5% (w/v) Pluronic F-127, 5 mg/mL BSA, and 0.2 mg/mL α-casein, and with 1 µL of appropriately diluted DDB complex. The mixture was incubated on ice for 30 minutes.

Following incubation, 40 µL of motility buffer supplemented with 2 mM ATP, 2 mM biotin, and a gloxy oxygen scavenger system was added to the bead–protein mixture. Two 20-µL aliquots of the resulting solution were then introduced into the flow chamber. The chamber was sealed with vacuum grease to prevent evaporation.

#### Data Acquisition

Optical tweezers experiments were performed at room temperature using a C-Trap system combining optical trapping with total internal reflection fluorescence (TIRF) and interference reflection microscopy (IRM) (C-Trap Edge, LUMICKS). Microtubules were visualized using IRM. Trap stiffness was adjusted to approximately 0.08–0.1 pN/nm.

Optical trapping data were acquired at a sampling rate of 20 MHz and digitally down-sampled by a factor of 256, resulting in a final sampling rate of 78.125 kHz. An anti-aliasing filter was applied to achieve a passband of 31.25 kHz. The resulting data were fitted to a Lorentzian power spectrum for subsequent analysis.

#### Data Analysis

Optical trapping data were processed using a custom MATLAB program.

### Total Internal Reflection Fluorescence (TIRF) Assay

#### Complex Assembly

Dynein (dimer), dynactin, and BicD2 (dimer) were mixed in the presence of 2 mg/mL BSA at a 1:1:1 molar ratio to a final concentration of 120 nM each. The mixture was incubated at 4 °C overnight to allow complex formation.

For experiments including Lis1, Lis1 was added to the preassembled DDB complex at a final DDB:Lis1 ratio of 1:5. For experiments examining different binding orders of dynein, dynein, dynactin, BicD2, and Lis1 were mixed at a 1:2:2:10 molar ratio, with a final concentration of 60 nM for the first dynein. This mixture was incubated overnight, after which an equal amount of the second dynein was added and incubated for 1 hour on ice.

For experiments using wild-type dynein and dynein tail, 0.3 µM dynein tail (labeled with TMR) was mixed with 0.6 µM wild-type dynein (labeled with Alexa Fluor 488). This dynein/dynein tail mixture was then combined with dynactin, BicD2, and Lis1 to a final molar ratio of 1:1:1:5. For experiments using mutant dynein and dynein tail, 0.6 µM dynein tail (labeled with Alexa Fluor 647) was mixed with 0.6 µM mutant dynein (labeled with Alexa Fluor 488). The resulting dynein mixture was then combined with dynactin, BicD2, and Lis1 to a final molar ratio of 1:1:1:5.

#### Sample Preparation

Microtubule polymerization and flow chamber preparation were performed as described for the optical tweezers assay, with the addition of 1 µg of either Cy5-labeled tubulin or HiLyte488-labeled tubulin to visualize microtubules. After microtubules were immobilized on the coverslip surface, the assembled complex was diluted in motility buffer containing 60 mM HEPES (pH 7.2), 75 mM KCl, 2 mM MgCl₂, 1 mM EGTA, 1 mM DTT, 2 mM ATP, 0.5% (w/v) Pluronic F-127, 10% glycerol, 2 mM biotin, 10 µM Taxol, 2 mM ATP, 5 mg/mL BSA, 1 mg/mL α-casein, and gloxy (oxygen scavenger system). The solution was introduced into the flow chamber, which was then sealed with vacuum grease to prevent evaporation.

#### Data Acquisition

Images were acquired using BioVis software (BioVision Technologies) with an acquisition time of 200 ms per frame. SNAP-Surface Alexa Fluor 488 was excited using a 488-nm laser, SNAP-Cell TMR-Star was excited using a 561-nm laser, and SNAP-Surface Alexa Fluor 647 was excited using a 640-nm laser.

#### Data Analysis

Kymographs were generated using Fiji^80^, and velocities were analyzed using a custom MATLAB program.

## Graphic Visualization

Molecular structures in the figures were visualized using UCSF ChimeraX^81^.

## Acknowledgments

C. A. Walker, L. Rao, X. Liu, and A. Gennerich were supported by National Institutes of Health (NIH) grant R01GM098469 and acknowledge use of the LUMICKS C-Trap, funded by NIH grant S10OD034445-01. X. Liu also received support from a Pilot Project Grant from the Rose F. Kennedy Intellectual and Developmental Disabilities Research Center at the Albert Einstein College of Medicine. L. Rao, A. Shatarupa, H. Sosa, and A. Gennerich were supported by NIH grant R01GM147332. A. Shatarupa and H. Sosa were additionally supported by NIH grant R01GM113164. J. Yang and K. Zhang were supported by NIH grant R35GM142959. Molecular graphics and analyses were performed using UCSF ChimeraX, developed at the University of California, San Francisco, with support from NIH grant R01GM129325 and the Office of Cyber Infrastructure and Computational Biology, National Institute of Allergy and Infectious Diseases.

## Author Contributions

C. A. Walker, L. Rao, and J. Yang performed the experiments. L. Rao, X. Liu and A. Shatarupa produced and purified proteins; C. A. Walker, L. Rao, J. Yang and F. Berger analyzed the experimental data; A. Gennerich designed the research; C. A. Walker, L. Rao and A. Gennerich wrote the manuscript. K. Zhang, H. Sosa and A. Gennerich secured funding.

